# Flexible representation of dynamics allows precise tuning to the environment

**DOI:** 10.64898/2026.04.26.720916

**Authors:** Raz Leib, Sae Franklin, David W. Franklin

## Abstract

To precisely control our movements, the sensorimotor control system builds representations of the external world. However, the neural encoding of these representations – whether we encode our control as a function of specific kinematic or kinetic variables – and how we might then use these representations to build up complex actions is still unknown. Each prior study has provided support for a different reference frame or proposed mixture models to explain the complex findings. Here, we propose a different framework in which the sensorimotor control system uses multiple representations, allowing it to flexibly tune its representations to task demands. We directly test this by showing that the representation of external forces is totally governed by the nature of the experienced forces. Participants form a global force representation based on the coordinate system that best explains the forces experienced at different arm postures. We suggest that when formalizing such global representation, the system decides on the representation by considering the level of uncertainty it has in each of the coordinate systems. This flexibility and redundancy in the representations allows the precise tuning to any specific task within our complex environment, explaining our expertise in a wide range of tasks.

**Significance Statement:** The coordinate representation used for the planning and control of movement has long been a fundamental question in neuroscience. While many studies have investigated the representation of external dynamics to answer this question, there exists no coherent view of the coordinate frame used for this representation. We show that this inconsistency arises because the nervous system has multiple representations, combining them depending on the experienced environmental forces. This flexibility in the representation of the environmental dynamics allows precise adaptation to the complex and changing dynamics of our world.

## Introduction

In many daily activities, such as manipulating objects, we experience forces that can affect our performance if not properly represented. For example, holding an umbrella on a windy day without considering the wind direction and strength can cause it to break under the strain of the wind. One key issue in understanding how our brain builds a representation of such dynamics is the choice of coordinate frame, since external dynamics could theoretically be represented in many different coordinate frames. Initial studies suggested that the brain represents forces in joint coordinates, which is based on using the shoulder, elbow, and wrist joint torques to represent force (Shadmehr and Mussa-Ivaldi 1994, Shadmehr and Moussavi 2000, Malfait, Shiller et al. 2002, Malfait, Gribble et al. 2005). Other studies supported representation in Cartesian, which is based on vectors in a coordinate system located externally to the body (Criscimagna-Hemminger, Donchin et al. 2003, Malfait and Ostry 2004, Burgess, Bareither et al. 2007) or an object based coordinate systems, in which forces belongs to the hand or a hand-held object using the orientation vector connecting the shoulder and the object or hand in space (Yeo, Wolpert et al. 2015, Franklin, Batchelor et al. 2016). While initial studies argued for encoding of specific coordinate systems, more recent studies have suggested that learning is not represented by any single coordinate system, but instead is a mixture representation of all three coordinate systems (Berniker, Franklin et al. 2014, Leib and Franklin 2025).

Common to these studies is that the force representation was examined using a generalization paradigm; participants adapted to and formed a representation of an external force in one location in the environment and were then examined to see how they generalized the learned forces at a second, unexplored location. However, as forces were only ever experienced in one location, the overall nature of forces remains ambiguous, explaining the inconsistent results regarding force representation. Indeed, the one study that provided explicit information about the force representation suggested that the sensorimotor system can adapt to at least two natural coordinate systems, but is impaired when the representation of the task does not correspond to a relevant coordinate system (Franklin, Batchelor et al. 2016). These results suggest that when we experience force in a single location in space, the nature of forces across the entire space remains ambiguous, thus forcing the brain to estimate the optimal force in unexplored areas by weighing different dynamics representations (Leib and Franklin 2025). Removing this ambiguity, such as experiencing forces in multiple areas, allows the brain to shift from optimal estimation of force representation to fitting the representation to the external forces.

While one study suggested the sensorimotor control system can adapt to two natural coordinate systems (Franklin, Batchelor et al. 2016), the conclusion was only based on the extent of adaptation with no measurement of the representation. Here, using a novel experimental design that allows us to directly measure the sensorimotor control systems representation, we directly test whether the experienced force environment will shape the brain’s representation of the environment. We provide the first direct evidence that the sensorimotor system can change the combination between the three coordinate systems based on the force profile of the environment. By removing force ambiguity in the environment, we can drive the motor system to represent the force in a manner that matches the nature of the tasks.

## Results

Here, by testing force generalization, we show that participants are able to tune their representation of dynamics in accordance with the force profile of the environment. We do so by having participants adapt to novel environmental dynamics in which we have removed the ambiguity in the environment by introducing force fields at two training locations in space. When presented with these environments, participants set their force representation so that it best aligned with the presented force profile. Moreover, we show that we can change the specific representation of the participant to a Cartesian, joint or object representation by simply presenting different profiles at the two training locations, while keeping the arm configuration and motion locations unchanged. To achieve this result, we used a haptic robot to create a virtual environment (Figure 1A) in which participants performed reaching movements (Figure 1B) while experiencing a velocity-dependent force field that varied with movement direction (Leib and Franklin 2025). For each movement direction, perpendicular forces that differ in magnitude and direction, i.e., clockwise or counterclockwise (Figure 1C), perturbed the hand from the original movement path. A convenient way to represent this force field is to assign positive and negative signs to the clockwise and counterclockwise directions, respectively. By doing so, the force pattern as a function of movement direction can be illustrated as a cosine wave (Figure 1D), allowing us to examine generalization as changes in the amplitude and phase of the predicted forces in new arm postures. In a series of experiments, we applied these force fields in two small areas of the total reachable environment, termed *training 1* and *training 2*. Critically, the specific forces applied in these training workspaces were calculated according to a single (experiments 1 and 2) or combination (experiment 3) of coordinate systems (Figure 1E). The specific form of the force field in the training 2 workspace removed the ambiguity regarding the nature of the force of the entire environment. When experiencing a force field only in one area, the motor system can determine the best way to represent the force (Leib and Franklin 2025). However, by introducing a second force field in a second location, set according to a specific representation, we motivated the system to represent force using that representation. To test if the motor system indeed changed the representation after simultaneously adapting to the force fields, we used a force generalization paradigm in which we tested the extrapolated force pattern at another spatial location in the environment (test workspace) using a force channel technique (Scheidt, Dingwell et al. 2001, Milner and Franklin 2005). In this test workspace, participants always moved inside a virtual mechanical channel that constrained their movement to a straight line and allowed them to successfully reach the target. While moving inside the channel, participants generated forces on the virtual walls, expressing their predictions of the force pattern they could encounter while moving in the new workspace. Using these forces, we calculated the level of force compensation participants exhibited as a ratio between the produced and unscaled ideal forces.

**Figure 1:**
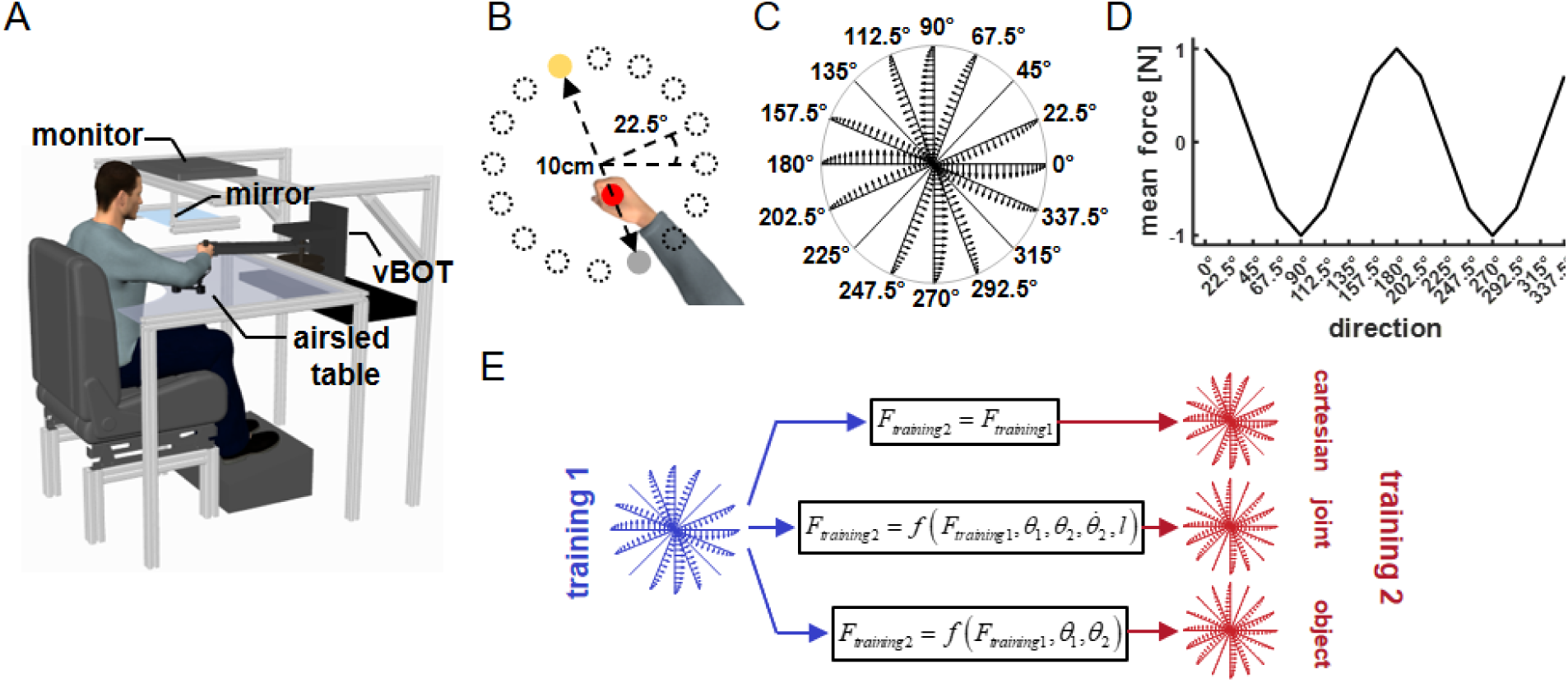
Experimental design. A) Participants sat in front of a robotic manipulandum (vBOT) while grasping the handle of the robot with their right hand. The participants’ arm was supported by an airsled, which reduced friction during movement. The virtual environment was projected on a mirror from a monitor mounted above the movement space. B) Display of the virtual workspace. The participant’s hand was represented by a red cursor. The task was to move from the start point (grey circle) to the target point (yellow circle). The start and target points were always located on the diameter of a 10 cm circle. There were 16 possible starting points evenly spaced along the circle’s circumference. On each trial, only one set of start and target points was presented. C) Endpoint force profiles as a function of movement direction. The forces were calculated using a scaled curl force field. The force profile that appeared in each direction is illustrated from the center to the boundary of the circle D) Cosine representation of the mean forces of the scaled force field as a function of movement direction. Clockwise forces were set as positive and counterclockwise forces were set as negative. E) Example for calculating the force profile used in training 2 workspace. We calculated the generalized force profile of the scaled force field using three different coordinate systems. For the Cartesian coordinate system, the force profile is identical to the scaled force field. For the joint and object coordinate systems, we need to use the arm configuration in the training workspaces, which results in a different force profile than the scaled force field. Changing the arm configuration changes these force profiles. In each experiment for training 2 workspace, we used a force profile calculated using one coordinate system or a combination of coordinate systems.

In Experiment 1, participants were assigned to one of three groups that differed in the force profile of training 2 workspace. All participants experienced the scaled force field in training 1 workspace (Figure 2A, blue force field) and one of three possible force fields in training 2 workspace, each calculated according to one of the three coordinate systems (Figure 2A, red force fields). The Cartesian-based force field was identical to the force profile in training 1 workspace (Figure 2 A2). The joint- and object-based force fields were similar to each other and were shifted compared to the scaled force field (Figure 2 A2, dashed and dotted red lines). In this case, the force profiles of these two reference frames were similar and differed from the Cartesian-based coordinate system. Thus, in this experiment, we could identify whether participants could change the weights between the Cartesian and the joint/object-based representations, but not between the joint- and object-based representations. That is, while participants in group 1 should prefer to represent the force using the Cartesian coordinate system, participants of groups 2 and 3, who experienced the shifted force field in training 2 workspace, could represent the forces using the joint and/or object coordinate systems. However, in the test workspace, the predicted generalized force pattern differed between all three coordinate systems. This allowed us to identify how participants represented the forces in the environment and, more importantly, whether the force profile in the training 2 workspace led them to change the weighting between Cartesian and joint/object-based coordinate systems.

**Figure 2:**
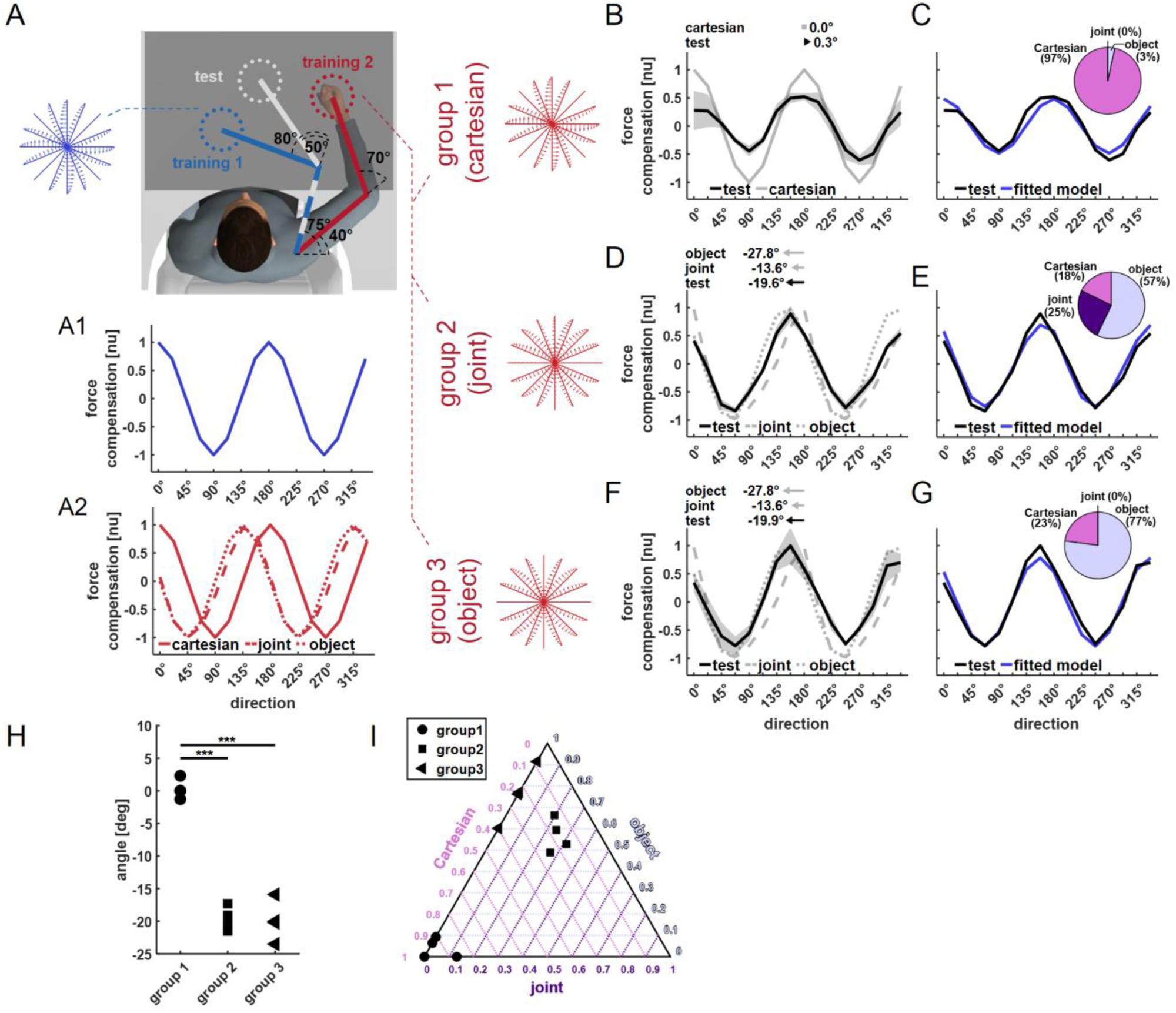
Experiment 1 demonstrates independent tuning of Cartesian compared to object/joint representations. A) Workspace locations and force fields used in Experiment 1. In training 1 workspace, all participants experienced the scaled force field (force field marked in blue). Participants were divided into three groups, each of which experienced a different generalized version of the scaled force field in training 2 workspace (force fields marked in red). These generalized versions of the scaled force field were calculated using the procedure described in Figure 1D using the three reference frames: Cartesian, joint and object. Summary force profiles as a function of movement direction for the force fields used in training 1 workspace (A1) and training 2 workspace (A2). Due to the arm configurations used in the training workspaces, the force profiles of the joint and object coordinate systems in the training 2 workspace were similar to each other but differed compared to the Cartesian-based force field. B) Mean force compensation curves exhibited at the test workspace for the Cartesian group (black curve, STD indicated using grey shaded area) and the predicted force generalization in the test workspace according to the Cartesian reference frame (grey curve). In this case, the Cartesian reference frame predicts that generalized forces will be identical to the scaled force field. Arrows above the panel indicate the shift of the mean force compensation profile and reference frame predictions relative to the original curve describing the trigonometrically scaled force field. C) Fitted force compensation curve for the Cartesian group based on a weighted sum between the Cartesian, joint, and object coordinate systems (blue curve). The weights between coordinate systems for this optimal fit were calculated based on minimizing the error between the mean experimental curve (black curve) and the fitted curve. The relative weight of each coordinate system is indicated in the inset pie chart. In this experiment and for this group, the weight of the Cartesian coordinate system was more dominant than that of the object and joint coordinate systems. D) Mean force compensation curves (as in B) at the test workspace for group 2, which had a joint-based force field in the training 2 workspace. In this case, since the force profile in the training 2 workspace did not differ between the joint and object coordinate systems, participants were expected to generalize the learned forces according to the joint (dashed grey line) or the object (dotted grey line) force representations. E) Fitted force compensation curve (as in C) for group 2 (joint). For this group, the object and joint weights were more dominant than the Cartesian coordinate system weight in the optimal fit. F) Mean force compensation curves (as in B) at the test workspace for group 3, which had an object-based force field in the training 2 workspace. G) Fitted force compensation curve (as in C) for group 3 (object). For this group, the fitted weights showed a significant contribution of the object compared to the Cartesian coordinate system, with zero weight for the joint coordinate system. H) Individual force compensation profile shift for each participant in groups 1-3. I) The relative weights between coordinate systems for the optimal fit, fitted to each participant’s force compensation profile in the test workspace. Each participant’s results are represented by a point in the Ternary plot. Each point represents the relative contribution of the Cartesian (pink, rightward diagonal), joint- (purple, leftward diagonal) and object-based (light blue, horizontal line) coordinate systems.

Participants in group 1, who experienced identical force fields in training 1 and 2 workspaces, exhibited a force compensation curve that resembled the cosine profile of the scaled force field (Figure 2B). The force compensation profile was not shifted compared with the scaled force field profile (0.3±1.5°, mean±STD). The profile and phase shift align with the prediction of a Cartesian-based force compensation profile, which predicts zero shift. In addition to the phase shift of the generalized force compensation pattern, there was also a reduction in the amplitude of the generalized forces. This aligns with previous results, which suggested that such a decrease in generalized force magnitude might be due to the distance between the training and test workspaces (Gandolfo, Mussa-Ivaldi et al. 1996, Berniker, Franklin et al. 2014). In addition to the phase analysis, we tested the relative contribution of each coordinate system when fitted to the experimental results. That is, we fitted a weighted sum of the generalized force compensation profiles based on the Cartesian, joint, and object coordinate systems. We determined the weight of each coordinate system in this mixture model to minimize the error between the experimental force compensation curve and the mixture model’s force compensation curve. The weight distribution of the mixture model showed that participants in group 1 relied mostly on the Cartesian representation (Figure 2C). When fitted to the data, we found that the optimal weights of this model were 0.47 for the Cartesian representation and 0 and 0.01 for the joint- and object-based representations, respectively. The relative contribution of the Cartesian coordinate system was 97%, suggesting that participants primarily represented this with a Cartesian force representation.

Switching the force field in training 2 workspace to a joint- or object-based force profile caused participants to change their force representation. For groups 2 and 3, we observed a shifted force compensation pattern when moving in the test workspace (Figure 2D and 2F). In both cases, the generalized force profile resembled a combination of the joint and object-based generalized force profiles. This was evident by the shift in the force compensation profiles for group 2 (-19.6±1.8°) and group 3 (-19.9±3.1°), which was between the predicted shift of the joint-based force profile (-13.6°) and the object-based force profile (-27.8°). When examining a fitted mixture model, we observed that in comparison with the results of group 1, for groups 2 and 3, the weight of the Cartesian coordinate system in the fitted mixture model decreased significantly (0.14 for group 2 and 0.19 for group 3) and instead the dominant weight was of the object-based coordinate system (0.47 and 0.66 for groups 2 and 3, respectively). For both groups 2 and 3, the combined relative contributions of the joint- and object-based representations made up 82% and 77% (Figure 2E and 2G). Importantly, although in this configuration we could not clearly induce participants to change the weights between the joint and object coordinate systems, the force compensation profiles of groups 2 and 3 differed from the force compensation curve of group 1, suggesting an ability to change the weight between a Cartesian force representation and a joint-/object-based representation.

Comparing the generalized curve shift results between the three groups (Figure 2H), we found the angle to be significantly different between the groups (F(2,9)=85.76 ,p<0.001, η^2^_p_=0.95). There was a significant shift of group 2 (t(9)=12.45, p<0.001, d=7.61) and group 3’s (t(9)=12.64, p<0.001, d=5.11) curve when compared with the curve of group 1. Since the force profile of workspace 2 was almost identical between groups 2 and 3, we did not find a significant difference in shift between these groups (t(9)=0.18, p=1.0, d=0.14). This supports the idea that removing force ambiguity by exposing participants to forces at multiple areas in the environment can change the way we represent forces, as evidenced by the change in generalized forces. Next, we aim to quantify the change in representation by comparing the fitted model results across the three experimental groups. Using Dirichlet regression, we found a significant effect of the experimental group on the relative contribution of each coordinate system (χ^2^=58.51, p<0.001). To compare between groups, we used the joint weight as a reference group (Maier 2014). As shown in Figure 2I, the contribution of the Cartesian coordinate system was the largest for group 1 and reduced significantly in group 2 (z=-8.12 ,p<0.001) and in group 3 (z=-2.12, p=0.03). Similarly, there was a significant increase in the relative contribution of the object-based coordinate system between groups 1 and 2 (z=10.38, p<0.001) and between groups 1 and 3 (z=11.03, p<0.001). In addition, there was a low contribution of the joint coordinate system across the three groups. This is a result of the similarity between the joint and object force profile used in the training 2 workspace, which triggered participants to rely more on the object coordinate system for force representation. Overall, the statistical analysis across the three groups suggests that participants adjusted the weighting between the Cartesian and the joint/object coordinate systems based on the force profile used in the training 2 workspace.

Following the change in force representation between the Cartesian and joint/object reference frames (Experiment 1), Experiment 2 tested whether force representation can change between the Cartesian/object and joint reference frames. Similar to Experiment 1, all participants experienced the scaled force field in training 1 workspace. The arm configuration used for the training workspaces was chosen to maintain the same hand orientation in space, thereby preserving the same force pattern in the object coordinate system (Figure 3A). In this case, the predicted force compensation profiles of both the Cartesian and object do not change between the training workspaces or in the test workspace. This sets two experimental groups: group 1 (Cartesian/object), which experienced the scaled force field in both training workspaces 1 and 2, and group 2 (joint), which experienced the scaled force field in training 1 workspace and a force field calculated according to the joint-based representation in training 2 workspace (Figure 3A1). For group 2, the force profile in the training 2 workspace was shifted in comparison to the scaled force field. In this case, participants in group 1 are predicted to generalize the force in the test workspace according to the Cartesian/object representation and exhibit the same profile of the scaled force field in the training and test workspaces. Participants in group 2 are predicted to represent the forces according to the joint-based representation and, in this case, exhibit a shifted version of the scaled force field in the test workspace. These predictions were confirmed as evident by the generalized force compensation curves in the test workspace for the two groups. Group 1 exhibited a generalized force profile similar to the scaled force field (Figure 3B), with a minor shift in the curve (2.2±2.4°). When we fitted the mixture model to the mean data (Figure 3C), we found that the weights of the Cartesian (0.32) and object (0.32) coordinate systems were more dominant than the joint coordinate system (0.18). Since the Cartesian and object representations were identical in this case, their combined relative contribution was 78% while the joint-based representation was equal to 22%. Group 2 exhibited a shifted generalized force profile (13.1±1.6°) that was similar to the predicted shift of the joint-based force compensation curve (13.7°, Figure 3D). Changing the force field in training 2 workspace also had a significant effect on the weights of the mixture model. The weights of the Cartesian and object-based coordinate systems greatly decreased (0.02 for each representation) while the weight of the joint-based coordinate system greatly increased (0.81). In this case, the joint-based representation had a large relative contribution to the model (94%) compared with the Cartesian/object representations (6%). Comparing the individual curve shift results between the two groups (Figure 3F), we found a significant increase in group 2 (t(6)=7.86, p<0.001, d=5.56). In addition, fitting the combined model to the individual data (Figure 3G), we again found a significant effect of the experimental group on the relative contribution of each coordinate (χ^2^=46.16, p<0.001). The relative contribution of the Cartesian/object coordinate system reduced while the contribution of the joint coordinate system increased between group 1 and group 2 (z=10.84, p<0.001). Overall, and similar to the results of Experiment 1, we could change the force representation by removing ambiguity from the environment. In this case, the coordinate system combination shifted from relying mainly on the Cartesian/object-based representations to relying mainly on the joint-based representation.

**Figure 3:**
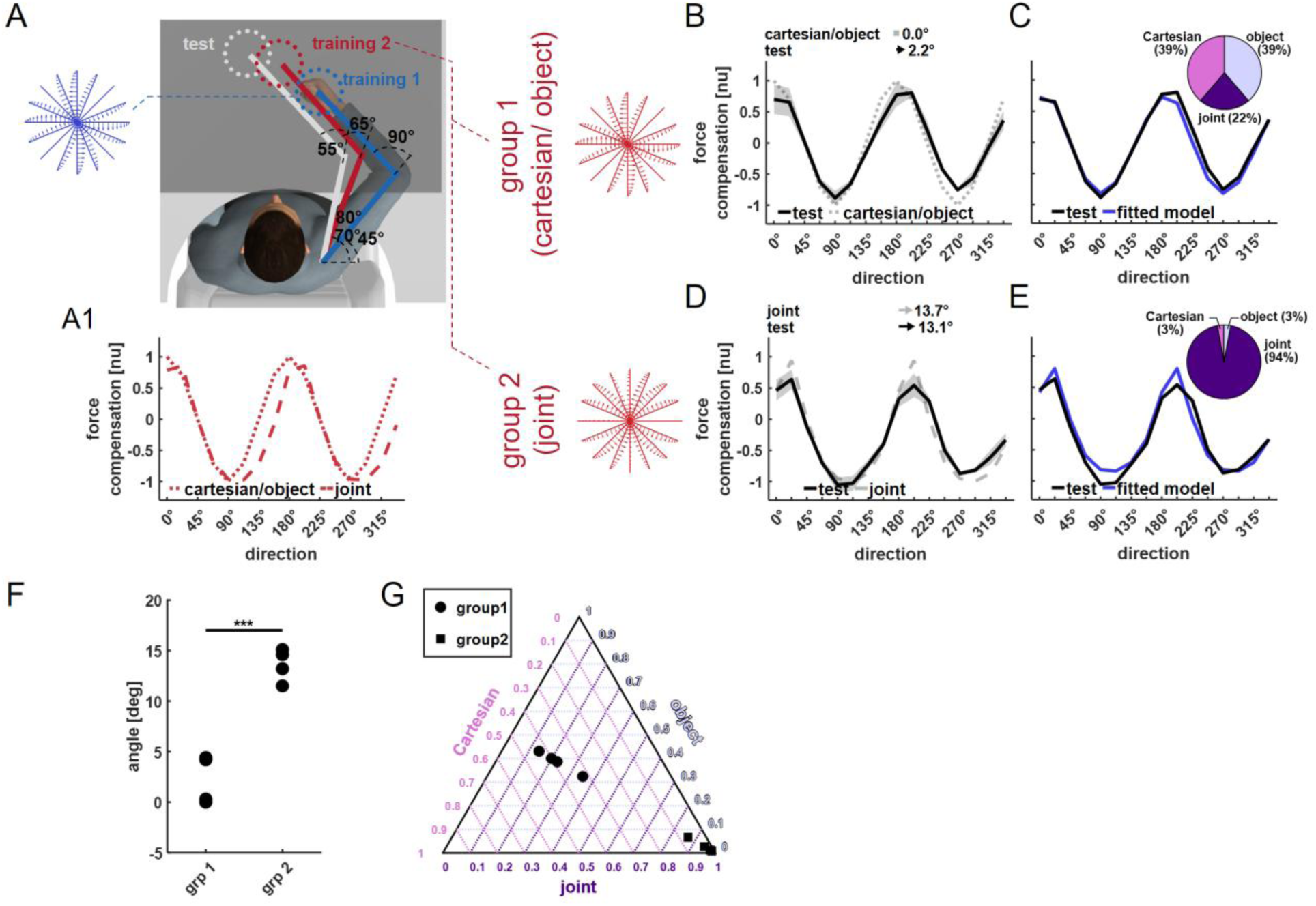
Experiment 2 demonstrates that the force representation can be tuned to either a Joint or Cartesian/object representation. A) Workspace locations and force fields used in Experiment 2. Similar to Experiment 1, both groups experienced the scaled force field in training 1 workspace (blue force field). In this experiment, since the Cartesian and object-based coordinate systems have the same force profile in training 1 and 2 workspaces, we did not have three experimental groups but only two: one group where the two force fields matched the Cartesian/object coordinate system, and a second group where the force fields matched according to the joint coordinate system. Panel A1 shows the force profiles as a function of movement direction for the force fields used in training 2 workspace. B) Mean force compensation curves exhibited in the test workspace for the Cartesian/object group (black curve) and the predicted force generalization in the test workspace according to these representations (dotted grey curve). Arrows above the panel indicate the shift of the mean force compensation profile and reference frame predictions with respect to the original curve describing the force field. C) Mean experimental force compensation curve (black curve) and fitted force compensation curve based on a weighted sum between the Cartesian, joint, and object coordinate systems (blue curve) for the Cartesian/object group. The relative weight of each coordinate system is indicated in the inset pie chart. In this experiment and for this group, the weight of the Cartesian/object coordinate system was more dominant than that of the joint coordinate systems. D) Mean force compensation curves at the test workspace for group 2 (as in B), which had a joint-based force field in the training 2 workspace (dashed grey line). In this case, since the force profile in the training 2 workspace did not differ between the joint and object coordinate systems, participants were expected to generalize the learned forces according to the joint (dashed grey line) or the object (dotted grey line) force representations. E) Fitted force compensation curve for group 2 (as in C). For this group, the weight for the joint-based coordinate system was more dominant than the weights of the Cartesian/object coordinate systems in the optimal fit. F) Individual force compensation profile shift for each participant in groups 1 and 2. G) Ternary plot of the relative Cartesian (pink, rightward diagonal), joint (purple, leftward diagonal) and object light blue, horizontal line) weights according to the optimal fit, fitted to each participant’s force compensation rofile in the test workspace.

Following Experiments 1 and 2, in which we demonstrated switching between force representations, we aimed to examine whether removing the environment ambiguity can allow for a force representation based on all three coordinate systems. In Experiment 3, we used the same arm configurations as Experiment 1 (Figure 4A). However, here we examined whether participants could generalize forces as an average of the three reference frames. That is, we introduced a force field in the training 2 workspace during the training, which was an average of the generalized force profiles for the Cartesian, joint and object coordinate systems in training 2 workspace (Figure 4A1). In this case, participants should exhibit a generalization pattern that differs from the observed generalization patterns of Experiment 1 and instead exhibit a generalization pattern similar to an average of the three generalization patterns. Indeed, we found that participants exhibited a generalization pattern that can be roughly predicted by averaging the three coordinate systems (Figure 4B). There was almost a perfect match in the shift angle between the predicted force compensation profile (-12.3°) and the experimental mean force compensation profile (-13.3±3.6°). Moreover, comparing this force profile with the force profiles of the three experimental groups that participated in Experiment 1, we observed that the force profile did not resemble any of the force profiles from Experiment 1, despite the identical arm configurations between experiments. This was also supported by a fitted mixture model that had a different set of weights in which the Cartesian and object coordinate systems were dominant (Fig. 3 C). In addition, we found that the fitted weights for individual force compensation data (Figure 4D) do not resemble the individual weights of any of the three groups in experiment 1. These weights were not significantly different from each other (Hotelling’s T² = 12.56, F(2,2)=4.18, p=0.073), which is consistent with the equal weights used to generate the training 2 workspace in this experiment.

**Figure 4:**
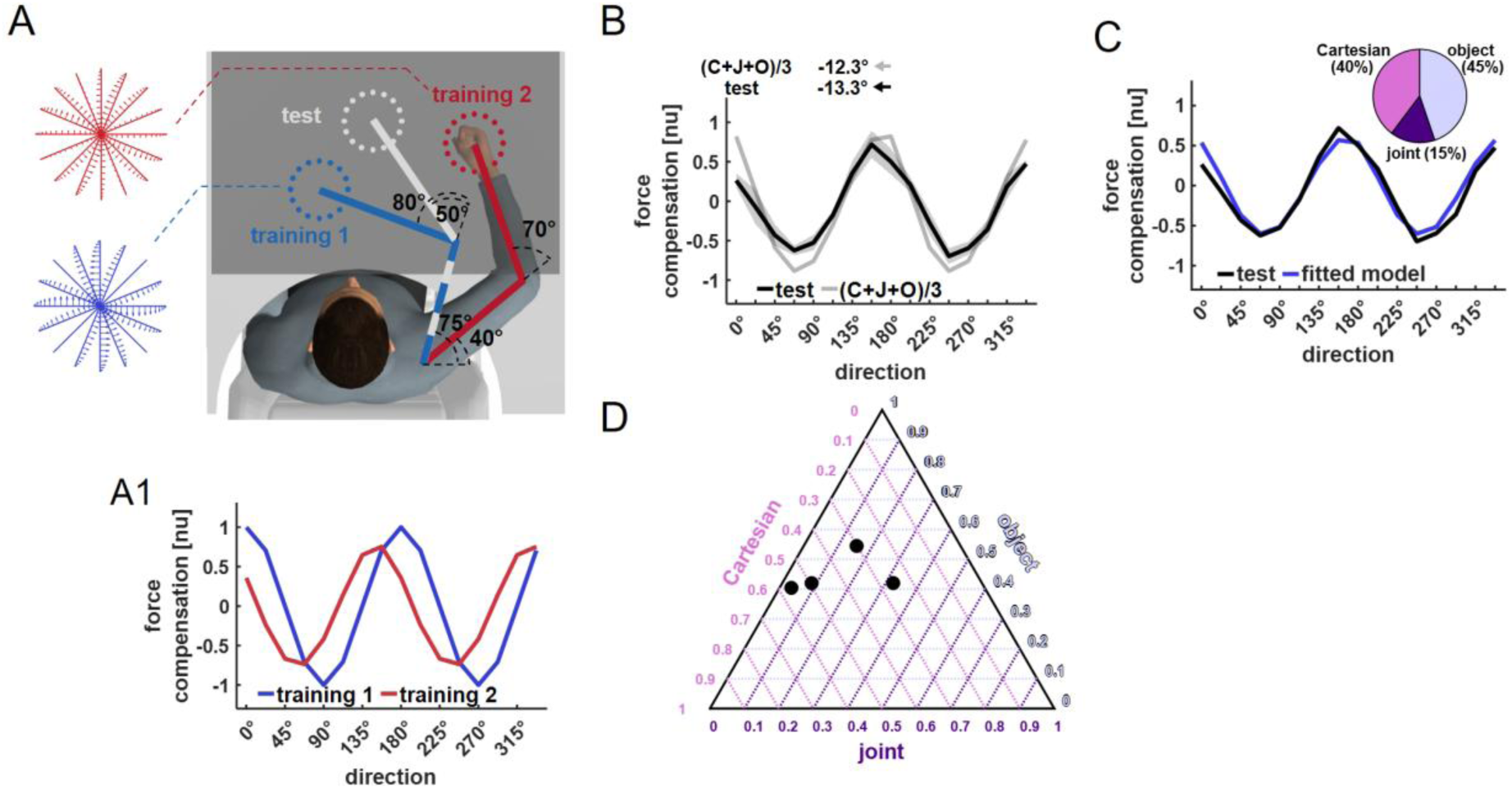
Experiment 3 demonstrates that the force representation can also be tuned to a mixture of coordinate systems. A) Workspace locations and force fields used in Experiment 3. These workspaces are identical to the workspaces used in Experiment 1. In training 1 workspace, participants adapted to the scaled force field. In training 2 workspace (panel A1), we used an average of the generalized force profiles of the three reference frames. That is, participants adapted to a force field that was the average of the signals in Figure 2A (bottom panel). Panel A1 represents the force profiles as a function of movement direction for the force fields used in training 1 workspace (blue curve) and training 2 workspace (red curve). B) Mean force compensation curves exhibited in the test workspace (black curve) and the predicted force generalization in the test workspace (grey curve). The predicted force was calculated as the average of the three reference frames. Arrows above the panel indicate the shift of the mean force compensation profile and reference frame predictions with respect to the original curve describing the force field. C) Mean experimental force compensation curve (black curve) and fitted force compensation curve based on a weighted sum between the Cartesian, joint and object coordinate systems (blue curve). The relative weight of each coordinate system is indicated in the inset pie chart. In this experiment, the weights of the Cartesian, joint and object coordinate systems were more balanced, demonstrating that force compensation relied on all three coordinate systems. Importantly, the weights differed from the fitted weights of Experiment 1 due to the different force profile used in training 2 workspace. D) Ternary plot of the relative Cartesian (pink, rightward diagonal), joint (purple, leftward diagonal) and object (light blue, horizontal line) weights according to the optimal fit, fitted to each participant’s force compensation profile in the test workspace. In this experiment, the individual data is close to the center of the triangle, indicating the fitted weights are roughly equal.

## Discussion

The ability to represent information and control movements in multiple ways can enable us to easily interact with different environments under many different conditions. For example, robotic systems that switch between force and position control can produce accurate movements in free space as well as during interaction with elastic objects (Raibert and Craig 1981). However, having such a capability also poses a scientific challenge in understanding the basic principles of the sensorimotor system (Franklin and Wolpert 2011, Leib, Howard et al. 2024). If the brain can switch between different controlled variables, contradicting evidence can be found even during simple operations such as a reaching movement. Here, we showed that during adaptation to a force field, the brain can change the combination between the Cartesian, joint and object-based coordinate systems used to represent dynamics. We triggered the change by removing environmental ambiguity regarding the global force nature. That is, when we experience forces that align with a specific representation, we can change our force representation to this representation type and generalize the force in new and unexplored areas in the environment according to it.

Our results suggest that the brain can switch between force representations, implying that the brain area responsible for force representation either holds multiple representations according to different coordinate systems and switches between them or changes the functionality of the neurons to represent one of the representations as needed. While we cannot differentiate between these two options, we can hypothesize that changes in task parameters altered the functionality of neurons. This is based on neural activity changes in the motor cortex (Li, Padoa-Schioppa et al. 2001) and the supplementary motor area (Padoa-Schioppa, Li et al. 2004) of monkeys adapting to a force field. Similarly, in other cases of representation redundancy, we can observe changes in the firing patterns of individual neurons. For example, in the motor cortex, neurons that exhibit a high firing rate at specific movement directions, i.e., preferred direction tuning, can change their tuning and shift it for different phases of the movement (Suway, Orellana et al. 2017) or as a function of where the movement is conducted in space (Caminiti, Johnson et al. 1991). Considering such examples, we can assume that the exposure to a second force field allowed the sensorimotor system to infer the representation that matches the experienced force pattern and change neurons’ functionality accordingly.

The process by which such a transition in force representation occurs is yet to be discovered. To evaluate such a process, we need to identify the base force representation that is used after adaptation to the force field in one location but before the exposure to the force field in the second location. In this case, the nature of the force across the entire environment is ambiguous, and the system can decide on the combination between different representations (Leib and Franklin 2025). Following adaptation in one location and quantifying generalization in other locations, it is necessary to remove the environmental ambiguity by introducing a second force field. However, the process of forming two separate force representations, one after the other, might be different from the process used in this study to form force representation based on an unambiguous environment. That is, it might be that the process of switching force representation after one was already formed can have different characteristics than forming an initial force representation.

Nonetheless, as we showed in our previous study, such a process might relate to measuring the reliability of different reference frames and combining them based on this reliability. Since noise creates an error between planned and executed force production, which depends on the arm state, the motor system will rely more on the representation for which the plan is less affected, reducing the uncertainty regarding the produced force (Leib and Franklin 2025). Removing the ambiguity of the environment by introducing a second force field serves this function precisely; the error between the planned force and the force profile at the second location vanishes almost entirely for one specific combination of reference frames, while this error increases for all other combinations. For example, if we are exposed to the same force profile in two locations, only by relying on a Cartesian-based coordinate system can we minimize the prediction error of the forces we will encounter when moving in each location. Relying on other coordinate systems or combinations between them will result in an elevated force prediction error, as they predict differences in the force pattern between the two locations due to the generalization nature of these coordinate systems. Thus, the sensorimotor system will choose to base its force representation on the Cartesian-based coordinate system, and as a result, we can also find that it generalizes the force according to this system (see Appendix).

While the idea of setting the weighting between coordinate systems based on error minimization predicts the generalized force compensation almost perfectly, there are still small discrepancies between the model and experimental results. Aside from the natural variability that may cause these discrepancies, prolonged exposure to the force fields might also tune the weights further. While the exposure to the force field in the experiment was extensive and allowed participants to form a representation of the force fields, longer interaction with the training force field might change parameters related to the force representation (Franklin, Nietschmann et al. 2025). Specifically, since we used a complex force field that varies in amplitude and direction with movement direction, having a well-tuned representation of the force profile for each direction can adjust the weights of each coordinate system. Future studies can examine how such long-term exposure to an unambiguous force field changes these weights.

We showed that the nature of the second force field set the coordinate system used for force representation. However, we cannot choose any force field for the second workspace due to the interference effect in forming multiple motor memories. Interference can occur when we attempt to adapt to multiple force fields simultaneously, when they appear randomly at the same spatial location, resulting in a lack of adaptation to either force field (Hwang, Donchin et al. 2003, Howard, Wolpert et al. 2013). Among the ways to avoid interference, we can use spatial differences in the locations where each force field is presented. That is, the spatial location of each force field serves as a contextual cue that allows the formation of different motor memories (Wainscott, Donchin et al. 2005, Howard, Ingram et al. 2012, Howard, Wolpert et al. 2013). However, if the difference in force patterns does not match the spatial difference in which they are introduced (Yeo, Wolpert et al. 2015) or the force patterns do not comply with any combination of valid coordinate systems (Franklin, Batchelor et al. 2016), the motor system would not be able to represent the environmental force in what we interpret as interference.

Our study shows both the plasticity and the ability to directly tune the force representation according to the nature of the force in the environment. Participants formed force representation according to a specific combination of coordinate systems when they were introduced to forces at multiple locations that represented the same coordinate systems. This supports the idea that the sensorimotor system employs multiple coordinate systems for force representation and utilizes a specific combination of representations to minimize the likelihood of misestimating forces in unexplored locations. Therefore, in ambiguous environments, the sensorimotor system represents forces to minimize internally induced force production errors, while removing the ambiguity allows it to tune the force representation to minimize external force errors.

## Methods

### Participants

Twenty-four right-handed participants (ten females; aged between 21-34) participated in one of three experiments (12 participants in experiment 1, 8 participants in experiment 2, and 4 participants in experiment 3) after signing a consent form and completing a handedness test (Oldfield 1971). The experimental protocol was approved by the Ethics Committee of the Medical Faculty of the Technical University of Munich. Each participant only performed one of the three experiments.

### Experimental Setup

We used the vBOT robotic device (Howard, Ingram et al. 2009) to create a haptic augmented virtual reality environment. The robotic device was used to record hand position and velocity while generating force feedback at 1 kHz in real-time. A six-axis force transducer (ATI Nano 25; ATI Industrial Automation) measured the end-point forces applied by the participant on the handle and was also sampled at 1 kHz. Participants were seated in front of the experimental setup and viewed a semi-silvered mirror that displayed the projection of an LCD screen placed directly above it (Figure 1A), allowing for visual feedback to be presented in the plane of arm movement. This mirror system prevented participants from viewing their hand. The participant grasped the robotic handle with their right hand, their arm supported on an air sled, allowing for frictionless movement. Once the participant was seated in front of the system, we measured the shoulder position relative to the origin of the virtual environment. To do so, we initially measured the upper arm length and the length between the elbow joint to the handle center. Next, we placed the hand at the system origin location by applying forces using the robot and measured the elbow and shoulder angles. Using these measurements and the inverse kinematics equations of a two-link arm model, we calculated the location of the shoulder.

Participants performed reaching movements from a start position (a 1.5 cm diameter gray circle) to a target position (a 1.5 cm diameter yellow circle) while viewing a cursor (a red circle with a diameter of 0.7 cm) representing their hand on the screen. The start and target positions were chosen from 16 equally spaced locations on a 10 cm diameter circle (Figure 1B). Each trial began with the robotic system positioning the hand at the start. After a random delay (0.75-1.5s), an auditory beep prompted movement towards the target. Participants were instructed to move with a consistent peak speed of 50cm/s. To do so, we presented visual feedback about the peak speed of their movement at the end of each movement. Speeds within ±8 cm/s were ’good’, slower speeds got a ’too slow’ message, faster speeds got a ’too fast’ message, and overshooting the target by more than 2 cm resulted in an ’overshoot’ message. For successful movement, we increased the score counter that appeared at the top of the screen by one point.

On each trial, the robot was used to produce one of four virtual environments. The first was a null field (NF), in which participants were free to move without experiencing any forces from the robot. For the other three environments, the robot applied forces on the participants’ hand during movement. The second environment was a *virtual mechanical channel*, on which we implemented two virtual walls that constrained the participants’ hand to move in a straight-line path to the target (Scheidt, Dingwell et al. 2001, Milner and Franklin 2005). The virtual walls were created by generating forces that were perpendicular to the line connecting the start and target positions. The forces were calculated according to the deviation of the hand position from the straight-line path multiplied by a stiffness coefficient of 4000 N/m and the velocity in the perpendicular direction multiplied by a damping coefficient of 2 Ns/m. The third type of virtual environment was the *force field 1*. During these trials, participants experienced forces acting on their hand that were a function of the hand’s instantaneous velocity. The forces were calculated using a scaled curl matrix

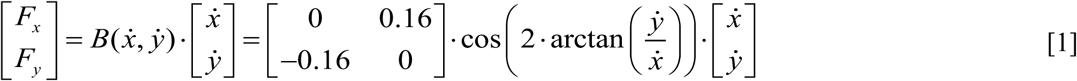

where *F_x_* and *F_y_* are the forces generated in the x and y axis, respectively and *x* and *y* are the hand velocities in the x and y axis, respectively. The traditional curl force field was scaled using a cosine trigonometric function of movement direction. That is, the magnitude and sign of the scaling factor changed according to movement direction, calculated using the inverse tangent function (Figure 1C). For example, when the motion was made only in the x-axis, i.e., horizontal right to left or vice versa, the scaling factor was equal to 1, or when the motion was made only in the y-axis, i.e., moving away or towards the body, the scaling factor was equal to (-1). We define clockwise forces as positive and counterclockwise forces as negative. Simulated mean forces that participants experienced can be illustrated as a cosine curve when plotted as a function of movement direction (Figure 1D). This force field was always applied at the training 1 workspace.

The fourth type of virtual environment was the *force field 2*. Similar to the force field 1, during these trials, participants experienced forces acting on their hand that were calculated using the hand’s instantaneous velocity. This force field was calculated as a generalized version of the force pattern of force field 1. That is, we simulated how forces of the trigonometric scaled force field would generalize in space according to different coordinate systems and used different combinations of these force fields for different experiments (Figure 1E). This force field was applied at the training 2 workspace. In the next section, we provide a detailed explanation of the nature of coordinate systems and how the scaled force field is generalized in space according to each of them.

### Coordinate Systems and Calculations of Generalization Patterns

We considered three different coordinate systems, using which the motor system could represent the learned dynamics. The first coordinate system we considered is the Cartesian coordinate system. For this representation, the forces are invariant with respect to the orientation of the arm. Therefore, we used the force pattern of force field 1 also for force field 2. Participants are expected also to exhibit similar forces to the learned forces for any other workspace in the environment [1], resulting in a similar cosine signal of generalized forces as a function of movement direction.

The second coordinate system we considered is the joint-based coordinate system (Shadmehr and Mussa-Ivaldi 1994, Malfait, Shiller et al. 2002). For this representation, the generalized force pattern of the scaled trigonometric force field depends on the arm configuration in space. Therefore, when moving between the workspaces, the forces of force field 2 differ from the forces of force field 1. That is, the force pattern of force field 2 would be a transformed version of the cosine function describing the forces of force field 1. Similarly, the generalized forces in other workspaces in the environment would also differ from the force fields 1 and 2. To calculate the generalized forces profile, we used a two-link arm model in which ***θ*** = [*θ_s_* ,*θ_e_* ] represents a vector of the shoulder and elbow joint values. We calculate the generalized forces between two workspaces, which we mark as training 1 workspace and training 2 workspace. That is, since we know the learned force pattern in training 1 workspace, i.e. the trigonometric scaled force field, we use it to calculate the generalized force pattern for the training 2 workspace. Initially, we need to derive the learned dynamics in the joint coordinate system and then derive the generalized forces. In joint representation, the learned forces are represented as joints torques using the Jacobian, which forms the relation between the two variables, ***τ*** = *J^T^* · ***F*** , where ***F*** = ⌈_⌊_*F* , *F* ⌉_⌋_ is a vector of the end-point forces, ***τ*** = [*τ _s_* ,*τ_e_* ]is a vector of joints torques, and *J_ws_*_1_ is the arm’s Jacobian matrix for the training 1 workspace.

Similarly, the hand velocities are represented using joint angular velocity and the Jacobian matrix, ***v*** = *J_ws_*_1_ ·***θ*** , where ***v*** = [*x*, *y*] is the hand velocity vector. Thus, using the end-point forces generated by the force field (equation 1) we derived the learned joint torques

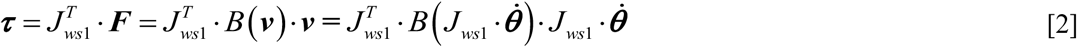

The expected forces during movement in other workspaces (training 2 workspace) can be computed using the learned torques and the arm’s Jacobian for the arm configuration at this workspace.

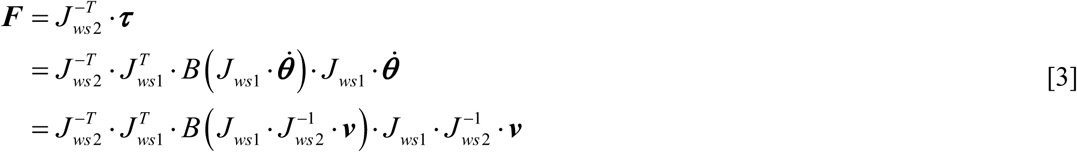

The third coordinate system we considered is the object-based coordinate system (Berniker, Franklin et al. 2014). For this representation, the motor system relates the dynamics to a grasped object. Therefore, the generalized forces are calculated according to the orientation of the held object, which is rotated with the hand orientation. For example, if we learned a force profile while moving straight away from the body and we now moved to another location in which the hand was rotated by an angle *θ* in external space, the learned profile is expected to appear in a direction that is rotated by *θ* . That is, initially, based on the velocity vector, we can find the learned force profile by rotating back to the original workspace ***F****_ws_*_1_ = *B*_(_*R*^−1^ (*θ* )· ***v***_)_· *R*^−1^ (*θ* )· ***v*** , where R is a rotation matrix. Then, we rotate these forces to the second workspace:

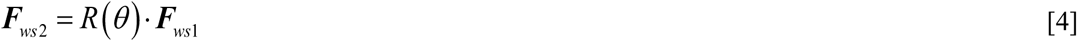

### Experimental Protocol

We conducted three experiments using similar experimental protocols (Figure 5). For each experiment, we used three workspaces (differing in their spatial location) in which participants performed the reaching movements; two training workspaces and one *test workspace*. Each experiment included four experimental phases. The experiment started with a familiarization phase of 16 trials that was performed only in the first training workspace (training 1 workspace). In this phase, participants performed reaching movements from each one of the sixteen starting points. The movements were unconstrained reaching movements in the null field, i.e. there were no forces applied to participants’ hand (null field trials). The second experimental phase, the pre-exposure phase, included recording baseline measurements of the movements in the force channel. For this purpose, participants performed a total of 384 movements in both the training and test workspaces. The movements included both unconstrained reaching movements and movements while the robot generated a virtual mechanical channel between start and target locations (channel trials). Participants performed 4 null field and 4 channel trials for each of the sixteen starting points in each workspace, resulting in 128 movements per workspace (a total of 384 movements across the three workspaces). For these trials, we randomized the start point and force field type using a uniform distribution. We divided the 384 into four blocks of trials and randomized the appearance of the condition, that is null field or channel trials, and the sixteen starting points for each of the three workspaces using a uniform distribution within a block. During the third, exposure phase, participants experienced the forces generated by the trigonometric-scaled curl force field (force field 1) in training 1 workspace and a generalized version of it (force field 2) in training 2 workspace. The coordinate system used to calculate the generalized force field 2 differed between experiments (see below). Participants performed 15 repetitions from each of the 16 starting points in each of the two training workspaces (a total of 480 trials). We divided the 480 trials into fifteen blocks, with each of the starting points appearing once. The order of appearance within a block was determined using a uniform distribution. Finally, after adaptation to the force fields, the fourth phase tested the generalization of the force field representation in the test workspace. During this phase, participants performed 1536 reaching movements in the training workspaces, where force fields were applied, and an additional 384 movements in the training and test workspaces, where their movements were constrained within the virtual mechanical channel. For each workspace, we had 8 repetitions of channel trials for each of the 16 starting points (a total of 384 trials). These trials were randomly introduced within the 1536 force field trials; for every series of 4 force field trials, we had one channel trial. The position of the channel trial within the series was random with a uniform distribution. Each participant participated in one of three experiments, and in the case of experiments 1 and 2, in one experimental group, which differed in the location of the training and test workspaces and the nature of force field 2.

**Figure 5:**
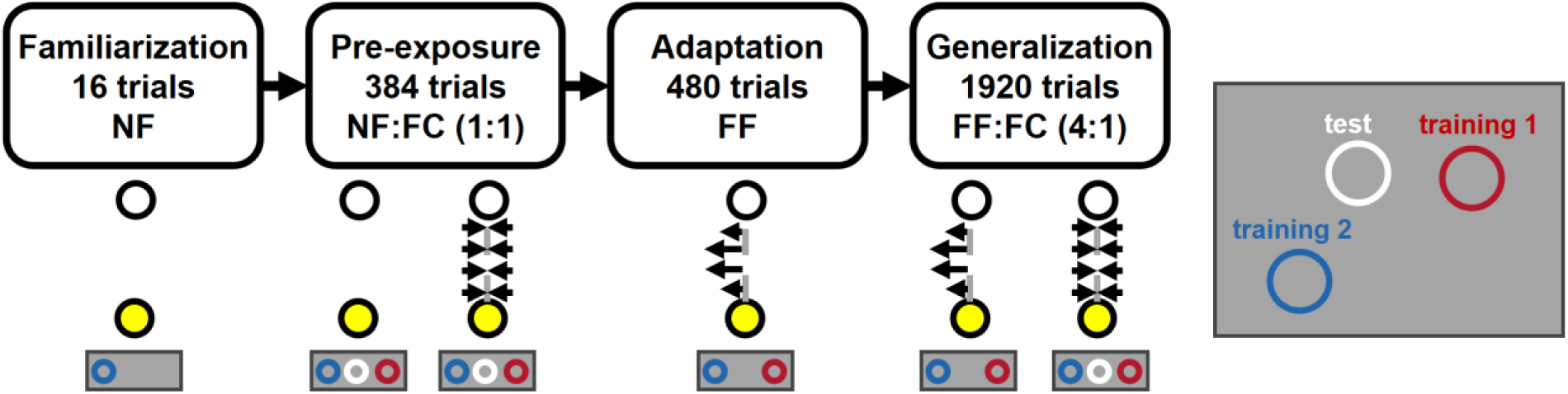
Experimental protocol. The initial familiarization phase included 16 trials in which no force was applied to the hand (NF condition). Movements at this phase were all done in the training 1 workspace. The pre-exposure phase included unconstrained movements (NF condition) and movements in a force channel (FC condition), with a ratio of 1:1 in all three workspaces. In the adaptation phase, participants performed the movements under the force field perturbation (FF condition) only in the training workspaces. In the last phase, we tested generalization using force channel (FC) trials introduced randomly in both the training and test workspaces, while participants kept moving under force field (FF) perturbations only in the training workspaces. The ratio of force field to force channel trials was 4:1.

*Experiment 1.* The location of the center of the training 1 workspace was set for each participant so that the participants’ shoulder joint was at 75° and the elbow joint was at 80°. At this workspace, we applied the force field 1 (the trigonometric scaled force field). The center of training 2 workspace was set so with participants’ shoulder angle at 40° and elbow angle at 70°. At this workspace, we applied a generalized version of force field 1. The center of the test workspace was set with the shoulder joint at 75° and the elbow at 50° (Figure 2A). In this experiment, we had three experimental groups that differed in the nature of force field 2. For group 1, the generalized force field was calculated using the Cartesian coordinate system. Similarly, for groups 2 and 3, the forces of force field 2 were calculated using the generalization of force field 1 based on the joint and object coordinate systems, respectively.

*Experiment 2.* This experiment was similar to experiment 1, but with different workspace locations (Figure 3A). The center of the training 1 workspace was set for each participant so that their shoulder joint was at 45° and the elbow joint was at 90°. The center of the training 2 workspace was set such that the shoulder and elbow joints were oriented at [70°, 65°]. The center of the test workspace was set such that the shoulder and elbow joints were oriented at [80°, 55°]. These workspace locations maintained a constant hand orientation while the arm configuration varied. Therefore, the generalization of force field 1 based on the Cartesian and object coordinate systems at training 2 and test workspaces was identical. As a result, we had two experimental groups that differed in the nature of force field 2. Group 1 had the generalized force field calculated based on the Cartesian/object coordinate system. For group 2, force field 2 was calculated by generalizing force field 1 using the joint coordinate system.

*Experiment 3*. In this experiment, we used the same arm configuration as in experiment 1 for the training and test workspaces. For the force field 2, we used a combination of all three generalization patterns according to the Cartesian, joint, and object coordinate systems. That is, we combined the generalization force pattern according to each coordinate system (Figure 2A) by summing them together, with each pattern getting a weight of 1/3: ***F*** = ^(^***^F^****^Cartesian^* ***^+ F^****^jo^*^int^ ***^+ F^****^object^* ^)^ . We used only one group of participants that adapted to the scaled force field in training 1 workspace and the average generalization force field in training 2 workspace.

In addition to calculating the force field 2 for each experiment and experimental group, we also simulated the expected forces in the test workspace according to the coordinate system that was used in each experimental group. For experiment 1 (Figure 2C and 2D), using a Cartesian based representation, the expected forces do not change when switching between workspaces. Thus, the force compensation profile would be similar to the profile of the force field for the test workspace. In the case of a joint-based representation, there is a leftward shift of the force compensation curve compared with the learned force field curve for some of the directions due to the change in arm posture. Similar to the joint-based representation, the force compensation profiles for object-based representation change as the hand’s orientation is changed between workspaces. For the test workspace, the curve was shifted to the left. Thus, leftward shift of the force compensation curve in the test workspace compared to the learned forces curve for groups 2 and 3 would suggest that the representation does not rely on a Cartesian based coordinate system. Similarly, no shift of the curve for group 1 would suggest that the representation relies more on a Cartesian based representation.

For experiment 2 (Figures 3C and 3D), the Cartesian and object-based representations predict that the expected force profile in the test workspace will be the same as the adapted force profile in the training workspaces. In the case of a joint-based representation, the prediction is of a shift to the right.

For experiment 3 (Figure 4C), the combination of the generalization patterns predicts a leftward shift of the force compensation curve in the test workspace since both the joint and object-based generalization patterns are shifted to the left (Figure 2C). However, here since we also combine it with the Cartesian based generalization force pattern, we expect the shift to be reduced compared with the joint or object-based generalization force pattern.

### Data Analysis

Data analysis was performed offline using Mathworks Matlab 2021a®.

*Force Compensation.* The force compensation was calculated during the generalization phase for the test workspace on the force channel trials. Force compensation in the test workspace demonstrates how the learned forces are generalized in space. Force compensation was calculated as the regression slope between the perfect force compensation signal, the independent variable, and the recorded force signal, the dependent variable (Smith, Ghazizadeh et al. 2006). The recorded force profile for each channel trial was identified between the start and end of the movement (first and last point at which velocity exceeded 5% maximum velocity). The perfect compensation force profile was determined as the force required for perfect compensation of movement in an unscaled curl force field and was calculated by multiplying the unscaled curl force field matrix by the velocity vector, [0 0.16 ; −0.16 0]·[*x*; *y*] . Here we compare the recorded force profile with forces computed using an unscaled version of the curl force (Leib and Franklin 2025). Once we calculated the force compensation measurement for each movement in the force channel, we averaged across the eight repetitions for each movement direction and repeated this for all 16 directions. The resulting pattern of the force compensation measure across all movement directions resembles a cosine function. To quantify phase differences between the scaled force field and the force compensation curve extracted from movements in the test workspace, we calculated the cross-correlation between these cosine signals. That is, the cross-correlation provided a measure for the shift of the force compensation curve from the designed scaled force field.

*Fitted mixture model.* We fitted a weighted sum of the generalized force compensation profiles based on the Cartesian, joint, and object coordinate systems. We determined the weight of each coordinate system in this mixture model to minimize the error between the experimental force compensation curve and the mixture model’s force compensation curve. We fitted the mixture model to individual force compensation curves, i.e., for each participant, and for the mean force compensation curve.

*Statistical Analysis.* To compare the differences in the shift of the generalized force compensation curve between groups of experiment 1, we used one-way repeated measures ANOVA with ‘group’ as a within-subject factor (3 levels) and the shift value as the dependent variable. We used post-hoc tests with a Bonferroni correction for multiple comparisons. For experiment 2, we compared shift differences between the two experimental groups using a pairwise t-test.

For experiments 1 and 2, we compared the fitted weights of a mixture model to the individual force compensation curves across experimental groups. Due to the compositional nature of weights, i.e., the weights as the dependent variables represent proportions that sum to one, we used Dirichlet regression. We used the groups as a categorical predictor. We implemented this analysis in R (v2026.07.1) using the ‘DirichletReg’ package (Maier 2014). For experiment 3, we tested whether the fitted weights differ between the three coordinate systems. To do so we first transformed the weights into open real space using the Isometric Log-Ratio (ILR) transformation. A one-sample Hotelling’s T² test was then conducted on the resulting two independent ILR coordinates to determine if the observed weight vectors significantly differed from a uniform, equal distribution (i.e., 1/3,1/3,1/3). The statistical analysis was performed in R using the ‘compositions’ package.

## Appendix

In this section, we calculated the Reliable Dynamics (Re-Dyn) model (Leib and Franklin 2025) predictions of the generalized force patterns across the three experiments. The Re-Dyn model calculates a weighted sum between the three reference frames (Cartesian, joint and object), considering the reliability of each force representation during dynamic generalization. That is, the model suggests that the motor system prefers certain representations if the execution of forces based on this representation is more robust to distortion arising from neural noise. In cases where forces are learnt in one workspace, the prediction of the weights is based on simulating the effect of internal neural noise on the variability in force production. In scenarios where we experience force fields in multiple, separate locations, the ambiguity regarding the overall nature of force is diminished, and thus the motor system will rely on a force representation that minimizes error in force production across the different locations. In this study, as we examined the latter case, we calculated the predicted weights of the Re-Dyn model by comparing the force predictions according to the three coordinate systems with the actual forces presented in the two training workspaces. That is, we first calculated the predicted force pattern in the training 2 workspace for each coordinate system based on the force pattern presented in the training 1 workspace. We included noisy arm state estimation when calculating the predicted force pattern to resemble the noise effect in the motor system (Leib and Franklin 2025). We then calculated the error in each movement direction for each force prediction and the pattern that was presented in the training 2 workspace. Since we presented a different force field in training 2 workspace for each experimental group in each experiment, we have different weightings between the three force representations. We repeated this simulation, varying the state noise (for more details, please see (Leib and Franklin 2025)) and calculated the variability of the error distribution. We then calculated the weight for each coordinate system based on the inverse-variance weighting. We used these weights to predict the force compensation profile in the test workspace and compared it with the experimental force compensation profile. This was repeated for each experiment and experimental group.

For Experiment 1, we found that the Re-Dyn model accurately predicted the shape and shift of the force compensation pattern exhibited in the test workspace (Appendix Figure 1). For experimental group 1 (Appendix Figure 1A, upper two panels), which had a Cartesian-based force field in training 2 workspace, we found that the Re-Dyn model predicted a shift of 0°, which matches the experimental results (0.3±1.5°, mean±STD). The predicted weights of this force compensation curve were mostly based on the Cartesian coordinate system, with some directions in which the weights of Cartesian and object-based representations were equal. For experimental groups 2 and 3 (Appendix Figure 1A, middle and bottom panels), which had a joint- and object-based force fields in training 2 workspace, we again found accurate prediction by the Re-Dyn model. The model could predict the leftward shift of the force compensation curve (for group 2, - 17.1° vs. -19.6±1.8° and for group 3, -25.5° vs. -19.9±3.1°) and the general shape of the curve. For these two groups, the joint- and object-based force representation had higher weight values compared with the Cartesian-based force representation.

The Re-Dyn model could also predict the results of Experiment 2 (Appendix Figure 1B). For experimental group 1, which experienced a force field based on the Cartesian/object coordinate system in training 2 workspace, we found that Cartesian/object-based weights were elevated compared to the joint-based weights. These weights generated a prediction that matched the experimental results (Appendix Figure 1B, upper two panels). The force compensation curve was predicted not to shift (0°), which aligns with the experimental curve that had a mean shift of 2.2° (±2.4° STD). Changing the force field in the training 2 workspace to a joint-based force field, the force compensation curve had a larger shift of 13.1° (±1.6°), which was similar to the Re-Dyn model prediction of a shift of 11.9° (Appendix Figure 1B, bottom two panels). This was due to the change in weights, which for this group were mainly based on the joint-based coordinate system.

When the force field in the training 2 workspace was calculated as a combination of the Cartesian, joint, and object coordinate systems, we found that the predicted weights for the three coordinate systems were similar to each other for most movement directions, and the difference between them was lower than in other experimental groups (Appendix Figure 1C, upper panel). Using these weights to produce the force compensation curve, we could predict the experimental result as both curves were similar to each other and had a similar leftward shift (-14.8° vs. -13.3±3.6°).

Across the three experiments, based on the error between the predicted force profile of each coordinate system and the force profile used in training 2 workspace, we were able to predict the force compensation profile. In each case, the weights of the coordinate system used in the training 2 workspace had elevated weight values, as it had the least error, which was primarily due to internal noise rather than a mismatch with the force profile used in the workspace. When we had a similar mismatch in force profile between coordinate systems (Experiment 3), the differences in weights were mainly because of noisy estimates of the arm state, as in the case of an ambiguous force field adaptation scenario (Leib and Franklin 2025).

**Appendix Figure 1:**
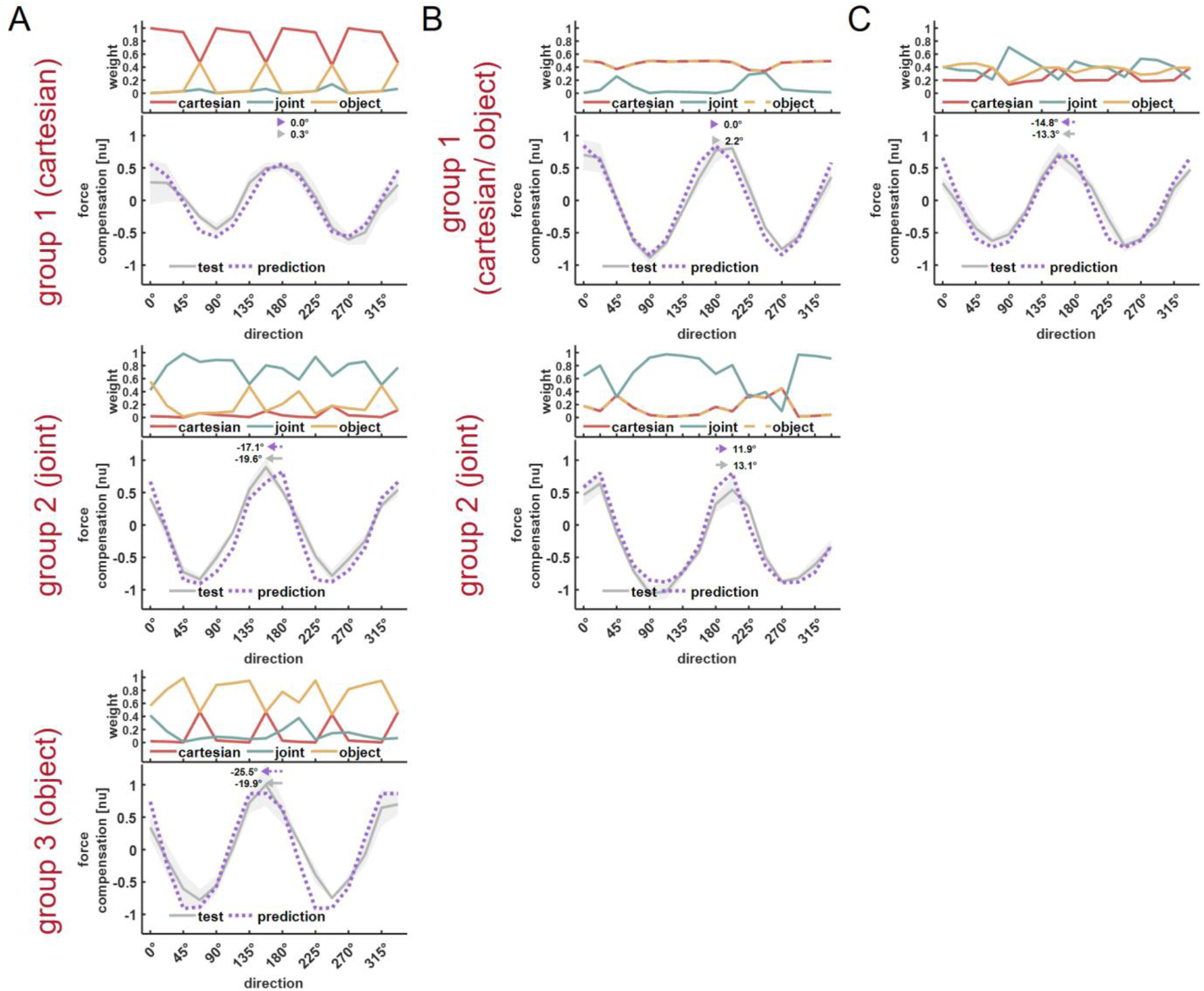
Re-Dyn model prediction for the generalization force pattern. A) Re-Dyn predictions for Experiment 1. For each experimental group, we calculated the prediction of the Re-Dyn model (dotted purple curve) and compared it with the experimental force compensation recorded in the test workspace (grey curve). To calculate the Re-Dyn predicted force compensation curve, we calculated a weighted sum between the Cartesian (red curve), joint (green curve), and object (yellow curve) force representations. Arrows above the force compensation curves indicate the shift of the mean force compensation profile and Re-Dyn prediction with respect to the curve describing the scaled force field (used in training 1 workspace). B and C) Re-Dyn predictions for Experiment 2. All notations are the same as in (A).

